# Concentration-dependent inhibition of *Streptococcus mutans* metabolic activity by purified *Croton lechleri* latex (sangre de grado) *in vitro*

**DOI:** 10.64898/2026.10.10.758117

**Authors:** Luis Fernando Rondán Villagra, Marco Zevallos Chávez, Jorge Franjo Rendulich, Marcelo Felix Ortega Suni, Rufo Alberto Figueroa Banda

## Abstract

**Background:** Chlorhexidine is the reference chemical antiplaque agent, but its adverse effects motivate the search for plant-derived alternatives. The proanthocyanidin-rich latex of *Croton lechleri* (sangre de grado) is a candidate, yet quantitative concentration–response data against *Streptococcus mutans* are scarce.

**Methods:** In this exploratory *in vitro* study, ethanol-purified *C. lechleri* latex was tested at working concentrations of 3–100% v/v (final in-well concentrations 0.6–20% v/v) against *S. mutans* ATCC 25175 in a 96-well resazurin assay with dual-wavelength reading (570/600 nm) and correction for medium and latex color. Chlorhexidine 1% (0.2% in-well) was the positive control. Each condition was run in technical triplicate within a single experiment. Data were analyzed by one-way ANOVA, Tukey’s test with simultaneous 95% confidence intervals (CI) and four-parameter logistic regression.

**Results:** Metabolic activity decreased monotonically with concentration, from 101.3 ± 1.7% (vehicle) to 54.0 ± 4.7% (12%),15.5 ± 4.0% (50%) and 8.6 ± 7.7% (100%), compared with 3.1 ± 2.8% for chlorhexidine (*F*_7,16_ = 160.7, *p* < 0.001). The lowest concentration that differed from the vehicle was 6% (difference 17.6 percentage points, 95% CI 2.1–33.1). The estimated half-maximal inhibitory concentration (IC_50_) was 12.3% v/v (95% CI 9.5–15.1). Differences from chlorhexidine at 50% and 100% were not statistically significant, but their confidence intervals were compatible with differences of up to 28 and 21 percentage points, respectively.

**Conclusions:** Purified *C. lechleri* latex inhibited the metabolic activity of planktonic *S. mutans* in a concentration-dependent manner. These preliminary findings do not establish equivalence with chlorhexidine and warrant replication in independent experiments with chemically standardized extracts expressed in mg/mL, determination of MIC/MBC and biofilm models.

**Resumen:** *Introducción:* La clorhexidina es el agente antiplaca de referencia, pero sus efectos adversos motivan la búsqueda de alternativas de origen vegetal. El látex de *Croton lechleri* (sangre de grado), rico en proantocianidinas, es un candidato, aunque faltan datos cuantitativos de concentración–respuesta frente a *Streptococcus mutans*.

*Métodos:* En este estudio exploratorio *in vitro* se evaluó el látex de *C. lechleri* purificado con etanol a concentraciones de trabajo de 3–100% v/v (0,6–20% v/v en el pozo) frente a *S. mutans* ATCC 25175 mediante resazurina en microplaca de 96 pozos, con lectura a 570/600 nm y corrección por medio y color del látex. La clorhexidina al 1% (0,2% en el pozo) fue el control positivo. Cada condición se evaluó por triplicado técnico en un único experimento. Se aplicó ANOVA de una vía, prueba de Tukey con intervalos de confianza (IC) simultáneos al 95% y regresión logística de cuatro parámetros.

*Resultados:* La actividad metabólica disminuyó de forma monótona con la concentración, de 101,3 ± 1,7% (vehículo) a 54,0 ± 4,7% (12%), 15,5 ± 4,0% (50%) y 8,6 ± 7,7% (100%), frente a 3,1 ± 2,8% con clorhexidina (*F*_7,16_ = 160,7; *p* < 0,001). La menor concentración que difirió del vehículo fue 6% (diferencia de 17,6 puntos porcentuales; IC 95% 2,1–33,1). La concentración inhibitoria media (CI_50_) estimada fue 12,3% v/v (IC 95% 9,5–15,1). Las diferencias frente a clorhexidina al 50% y 100% no fueron estadísticamente significativas, pero sus intervalos de confianza fueron compatibles con diferencias de hasta 28 y 21 puntos porcentuales. **Conclusiones**. El látex purificado de *C. lechleri* inhibió la actividad metabólica de *S. mutans* planctónico de forma dependiente de la concentración. Estos hallazgos preliminares no demuestran equivalencia con la clorhexidina y justifican su replicación con experimentos independientes, extractos químicamente estandarizados en mg/mL, determinación de CMI/CMB y modelos de biopelícula. **Palabras clave** *Croton lechleri*; sangre de grado; *Streptococcus mutans*; caries dental; resazurina; proantocianidinas

## 1 Introduction

Dental caries remains the most prevalent oral condition worldwide. In 2017, an estimated 2.3 billion people had untreated caries in permanent teeth [1], and untreated caries continues to lead the global burden of oral diseases [2]. Low-cost preventive adjuncts are therefore of particular interest in settings with limited resources.

*Streptococcus mutans* is a key cariogenic organism, not because it acts in isolation but because its glucosyltransferases (Gtfs) synthesize the extracellular polysaccharides that build the biofilm matrix. This matrix promotes adhesion, retains acids and creates the acidic microenvironments that drive enamel demineralization [3, 4].

Chlorhexidine is the reference chemical antiplaque agent and provides a clear adjunctive benefit on plaque and gingival inflammation, although tooth staining and taste alteration limit its long-term use [5]. Plant polyphenols, and proanthocyanidins in particular, inhibit Gtf activity, reduce *S. mutans* biofilm formation and attenuate caries development *in vivo* [6], which makes proanthocyanidin-rich natural products attractive candidates.

*Croton lechleri* Müll. Arg. (Euphorbiaceae) produces a red latex, known as sangre de grado or dragon’s blood, that is widely used in Amazonian traditional medicine [7]. Proanthocyanidins are its major constituents and can reach up to 90% of the dry weight of the resin [8]. A purified proanthocyanidin oligomer from this species, crofelemer, has reached clinical use as an antisecretory antidiarrheal agent [9]. Evidence against *S. mutans* is nevertheless limited. A disk-diffusion study reported inhibition zones with undiluted and 75% latex but not with 50% latex, all considerably smaller than those of 0.12% chlorhexidine [10], and other reports summarized in a recent review found no activity against *Streptococcus* sp. [8]. Agar diffusion provides largely qualitative information, depends on how readily the test material diffuses through agar and does not describe a concentration–response relationship.

Microplate assays based on the redox indicator resazurin provide a quantitative readout of bacterial metabolic activity and are widely used to screen phytochemicals [11, 12]. Because colored extracts can interfere with absorbance readings, they require appropriate optical controls. Here we report an exploratory microplate study of purified *C. lechleri* latex against *S. mutans* ATCC 25175. Our aims were to describe the concentration–response relationship, to estimate the half-maximal inhibitory concentration and to compare the observed effects with chlorhexidine using effect sizes and confidence intervals rather than *p* values alone.

## 2 Materials and methods

### 2.1 Study design

This was an exploratory, quantitative *in vitro* experimental study. All conditions were tested in technical triplicate (three wells per condition) within a single experiment. No sample size calculation was performed *a priori*, and the results are therefore presented as preliminary estimates with their confidence intervals. The study involved no human participants or animals. The workflow is summarized in Figure 1.

**Figure 1.**
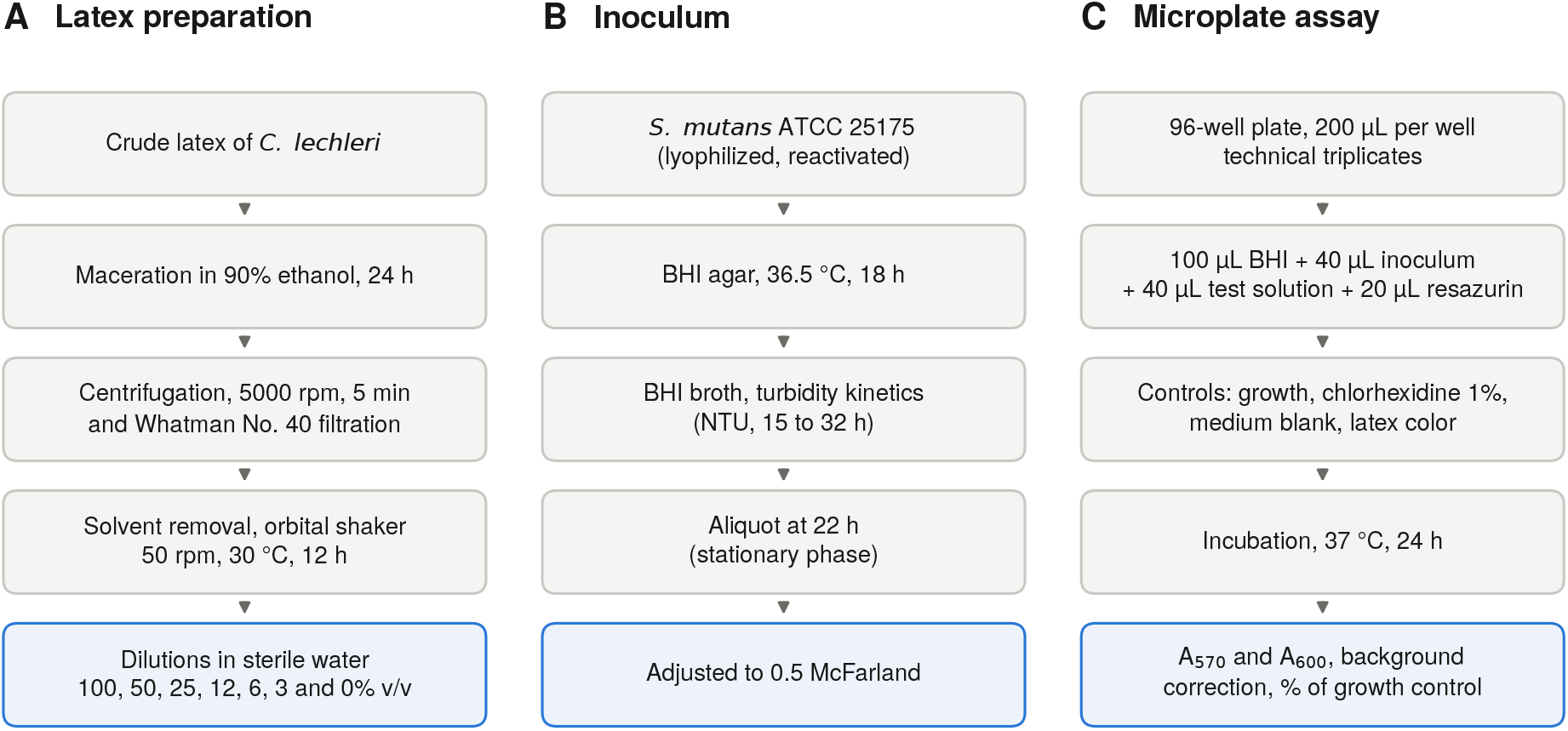
Experimental workflow. **(A)** Purification and dilution of *Croton lechleri* latex. **(B)** Preparation and standardization of the *Streptococcus mutans* ATCC 25175 inoculum. **(C)** Microplate resazurin assay. Highlighted boxes indicate the materials and the final readout that enter the assay.

### 2.2 Latex preparation

Crude *C. lechleri* latex was purified before use to remove insoluble impurities. The latex was macerated in 90% ethanol for 24 h and centrifuged at 5000 rpm for 5 min. The supernatant was recovered, the centrifugation was repeated and the supernatant was filtered through Whatman No. 40 paper. The solvent was then removed by evaporation on an orbital shaker (50 rpm, 30 °C, 12 h). The resulting purified latex (designated 100%) was diluted in sterile distilled water to working solutions of 50, 25, 12, 6 and 3% v/v. Sterile distilled water (0%) served as the vehicle control. Chlorhexidine 1% was prepared on the day of the assay by diluting a commercial concentrate in sterile distilled water, protected from light and kept at 4 °C until use (≤ 24 h).

### 2.3 Bacterial strain and inoculum

*Streptococcus mutans* ATCC 25175 was obtained in lyophilized form and reactivated for 48 h at 37 °C. Colonies grown on brain heart infusion (BHI) agar (Millipore-Merck, ref. 1104930500) at 36.5 °C for 18 h were transferred to BHI broth in 25 mL tubes, and growth was followed by turbidimetry (nephelometric turbidity units, NTU). Aliquots for the assay were taken at 22 h, when turbidity had reached a plateau (Table 1). Viable counts were not determined; approximate cell densities were estimated from a published relationship of 90 ± 5 NTU ≈ 1.5 × 10^8^ CFU/mL. The suspension was adjusted to 0.5 on the McFarland scale before dispensing.

**Table 1.**
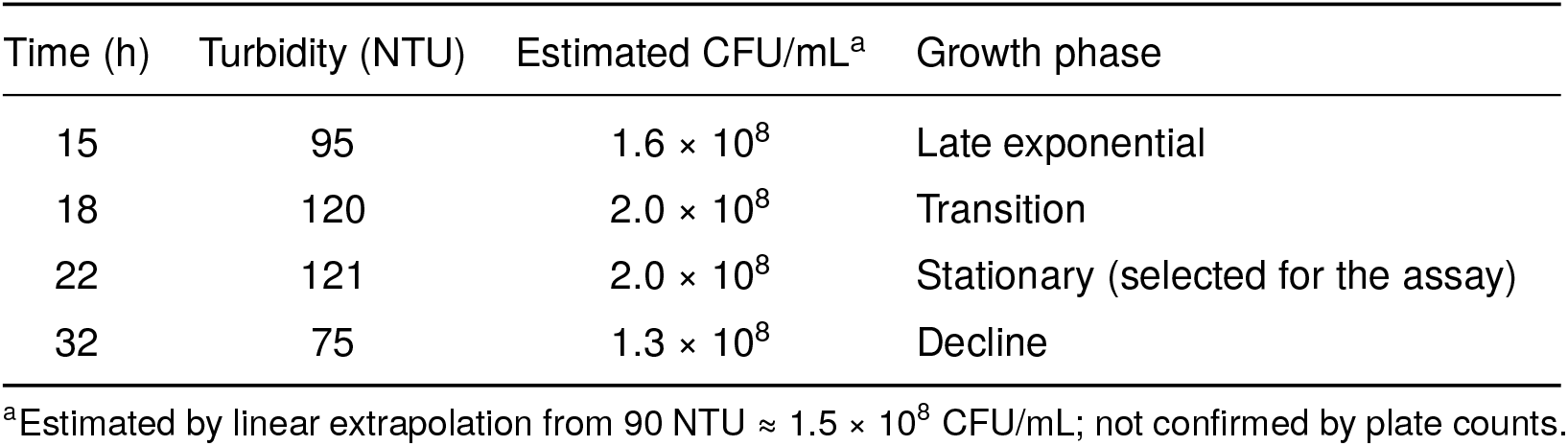
Turbidimetric kinetics of the *S. mutans* ATCC 25175 culture used to select the inoculum.

### 2.4 Microplate assay

Assays were set up in sterile 96-well plates in a biosafety cabinet with a final volume of 200 µL per well: 100 µL of BHI broth, 40 µL of standardized inoculum, 40 µL of latex working solution (or control) and 20 µL of 1% resazurin in distilled water. Each test solution was therefore diluted five-fold in the well, so that the 100, 50, 25, 12, 6 and 3% working solutions correspond to final concentrations of 20, 10, 5, 2.4, 1.2 and 0.6% v/v, and chlorhexidine 1% corresponds to 0.2% in the well. Controls in separate columns were (i) growth control (BHI and inoculum without treatment); (ii) positive control (chlorhexidine 1%, BHI and inoculum); (iii) medium blank (BHI without inoculum); and (iv) latex color controls (latex working solution, BHI and resazurin without inoculum), which correct for the color of the latex and for any reduction of resazurin not attributable to bacteria. Plates were mixed by gentle tapping, covered and incubated at 37 °C for 24 h.

### 2.5 Absorbance reading and outcome

After incubation, plates were gently homogenized, protected from light and read in absorbance mode at 570 and 600 nm. The corrected signal of each well was calculated as

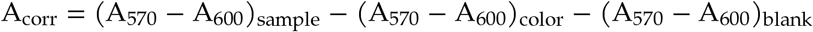

and metabolic activity was expressed as a percentage of the background-corrected growth control (100 × A_corr_/A_corr, growth control_). Inhibition was calculated as 100 minus metabolic activity. Readings outside the linear range of the instrument were discarded and repeated after a brief rest protected from light.

### 2.6 Statistical analysis

Results are presented as mean ± standard deviation (SD) with *t*-based 95% CI (*n* = 3 wells per condition). Groups were compared by one-way ANOVA on metabolic activity. Homogeneity of variances was assessed with Levene’s test. Because normality tests have very little power with three observations per group, normality was assessed on the pooled ANOVA residuals (Shapiro–Wilk). Pairwise comparisons used Tukey’s honestly significant difference test, and effect sizes are reported as mean differences in percentage points with simultaneous 95% CI; η^2^ is reported for the global effect. A four-parameter logistic model was fitted to the replicate-level data of the latex-treated wells (log-scaled concentration) to estimate the IC_50_, with asymptotic 95% CI and a residual bootstrap (2000 resamples) as a sensitivity check. A non-significant difference was not interpreted as evidence of equivalence. The significance level was 0.05. Analyses were performed in Python 3 with SciPy 1.18. Replicate-level data are provided in Supplementary Table S1.

## 3 Results

### 3.1 Concentration-dependent inhibition

Metabolic activity of *S. mutans* decreased monotonically as the latex concentration increased (Table 2; Figure 2A). Mean activity fell from 101.3 ± 1.7% with the vehicle to 93.8 ± 5.1% at 3%, 83.8 ± 6.2% at 6%, 54.0 ± 4.7% at 12%, 27.1 ± 8.4% at 25%, 15.5 ± 4.0% at 50% and 8.6 ± 7.7% at 100%. Chlorhexidine reduced activity to 3.1 ± 2.8%. The global effect was significant (*F*_7,16_ = 160.7, *p* < 0.001, η^2^ = 0.99), with homogeneous variances (Levene *p* = 0.95) and no evidence against normality of the residuals (Shapiro–Wilk *p* = 0.09).

**Table 2.** Metabolic activity of *S. mutans* ATCC 25175 after 24 h of exposure to purified *C. lechleri* latex and controls (*n* = 3 technical replicates per condition).

| Condition | In-well (% v/v) | Activity, % (mean ± SD) | 95% CI<sup>a</sup> | Inhibition, % | Group<sup>b</sup> |
| --- | --- | --- | --- | --- | --- |
| Vehicle (0%) | – | 101.3 ± 1.7 | 97.2 to 105.5 | –1.3 | a |
| Latex 3% | 0.6 | 93.8 ± 5.1 | 81.2 to 106.4 | 6.2 | ab |
| Latex 6% | 1.2 | 83.8 ± 6.2 | 68.4 to 99.1 | 16.2 | b |
| Latex 12% | 2.4 | 54.0 ± 4.7 | 42.4 to 65.5 | 46.0 | c |
| Latex 25% | 5 | 27.1 ± 8.4 | 6.3 to 47.8 | 72.9 | d |
| Latex 50% | 10 | 15.5 ± 4.0 | 5.4 to 25.5 | 84.5 | de |
| Latex 100% | 20 | 8.6 ± 7.7 | –10.5 to 27.7 | 91.4 | e |
| Chlorhexidine 1% | 0.2 | 3.1 ± 2.8 | –3.8 to 10.0 | 96.9 | e |

Activity is expressed relative to the growth control. Latex concentrations refer to the working solutions; the in-well column gives the final concentration after five-fold dilution. <sup>a</sup>*t*-based intervals with two degrees of freedom; negative lower limits reflect the small sample and have no physical meaning. <sup>b</sup>Conditions sharing a letter did not differ significantly (Tukey, $\alpha = 0.05$ ).

**Figure 2.**
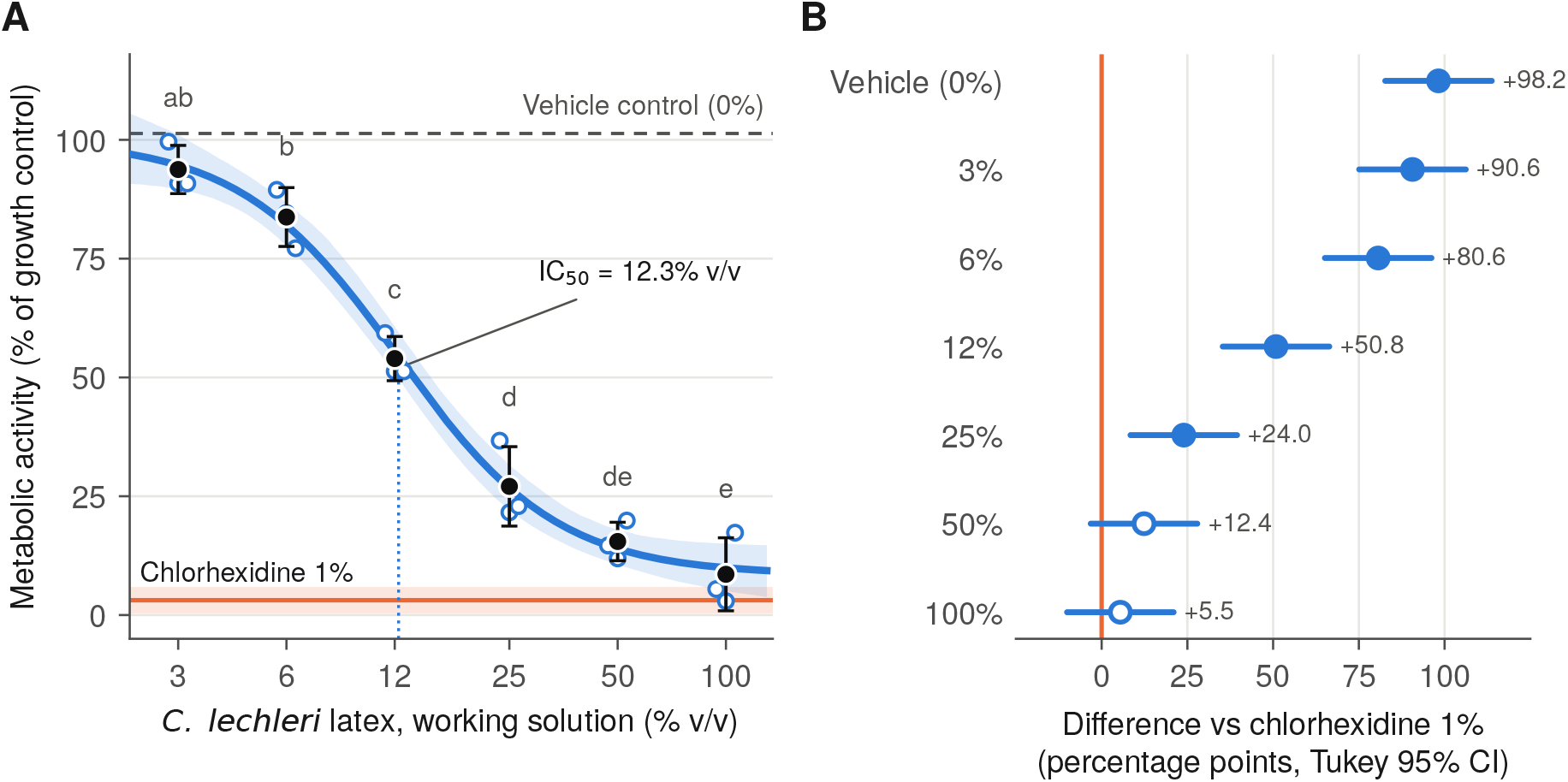
Effect of purified *Croton lechleri* latex on *Streptococcus mutans* ATCC 25175. **(A)** Concentration–response curve. Open circles are individual wells, black points are means ± SD (*n* = 3), and the blue line and band are the four-parameter logistic fit and its 95% bootstrap confidence band. The orange line and band show chlorhexidine 1% (mean ± SD) and the dashed line the vehicle control. Letters indicate Tukey groupings. Concentrations refer to working solutions (final in-well concentration is one-fifth). **(B)** Mean difference in metabolic activity between each condition and chlorhexidine 1%, with Tukey simultaneous 95% CI. Filled markers, *p* < 0.05; open markers, *p* ≥ 0.05. Positive values indicate less inhibition than chlorhexidine.

The lowest concentration that differed from the vehicle was 6% (difference 17.6 percentage points, 95% CI 2.1–33.1; *p* = 0.021), whereas 3% did not (7.6 points, 95% CI −7.9 to 23.1; *p* = 0.69). Each further doubling from 6% to 25% produced a significant additional reduction, while 25% and 50% and, likewise, 50% and 100% did not differ from each other (Table 2; Supplementary Table S2). The logistic model described the data well (*R*^2^ = 0.975) and estimated an IC_50_ of 12.3% v/v (95% CI 9.5–15.1; bootstrap 9.7–14.7), equivalent to approximately 2.5% v/v in the well, with a Hill slope of 2.0 and a lower plateau of 8.5% residual activity.

### 3.2 Comparison with chlorhexidine

All latex concentrations from 3% to 25% were significantly less inhibitory than chlorhexidine (Figure 2B). At 25%, residual activity exceeded that of chlorhexidine by 24.0 percentage points (95% CI 8.4–39.5; *p* = 0.001). The differences at 50% (12.4 points, 95% CI −3.2 to 27.9; *p* = 0.18) and 100% (5.5 points, 95% CI −10.1 to 21.0; *p* = 0.92) were not statistically significant. However, their confidence intervals were wide and remained compatible with substantially lower efficacy than chlorhexidine, so these results do not support a claim of comparable activity.

## 4 Discussion

In this exploratory study, purified *C. lechleri* latex reduced the metabolic activity of planktonic *S. mutans* ATCC 25175 in a concentration-dependent manner. The effect was detectable from the 6% working solution, was half-maximal at about 12% (approximately 2.5% v/v in the well) and approached near-complete inhibition at 50–100%. The steep, sigmoidal profile and the small residual activity at the highest concentrations indicate a consistent concentration–response relationship within this assay.

These findings agree in direction with the only previous study on *S. mutans* that we identified, in which undiluted and 75% latex produced inhibition zones that were smaller than those of chlorhexidine [10]. Our assay detected activity at considerably lower concentrations. Several differences may explain this, including the test format (broth microplate versus agar diffusion, in which large polyphenolic oligomers may diffuse poorly), the strain (ATCC 25175 versus ATCC 35668), the latex preparation (ethanol-purified versus crude) and the endpoint (metabolic activity versus zone diameter). Reports of no activity against *Streptococcus* sp. [8] further suggest that results depend on the source and preparation of the latex and on the method. This variability underlines the need for chemical standardization before results from different studies can be compared.

At the two highest concentrations, the difference from chlorhexidine was not statistically significant. This should not be read as equivalence. With three wells per condition, the confidence intervals for these comparisons included differences of up to 28 (50%) and 21 (100%) percentage points in residual activity. Demonstrating comparable efficacy would require a pre-specified equivalence or non-inferiority margin and an adequately powered design with independent replicates.

The proanthocyanidins that dominate the latex [8] are plausible contributors to the observed activity, given that proanthocyanidins from other sources inhibit Gtfs and reduce *S. mutans* biofilm formation and caries [6]. However, this study did not measure Gtf activity, glucan synthesis or biofilm formation, nor did it identify the active constituents. These mechanisms therefore remain hypotheses to be tested, not conclusions supported by the present data.

Some methodological aspects deserve comment. Resazurin reduction reflects metabolic activity rather than viable cell counts, so the assay cannot distinguish bacteriostatic from bactericidal effects. Polyphenol-rich extracts may also absorb in the visible range or reduce resazurin chemically. The latex color controls, containing latex, medium and resazurin without bacteria, were designed to subtract both effects, but they cannot exclude interactions that occur only in the presence of bacteria. Finally, the in-well inoculum (approximately 3 × 10^7^ CFU/mL, estimated) and its stationary-phase origin differ from standardized susceptibility protocols, which use about 5 × 10^5^ CFU/mL [13]. The assay was therefore more demanding than a standard test, and the values reported here are not minimum inhibitory concentrations.

### 4.1 Limitations

The results should be interpreted in light of the following limitations, each of which may bias the estimates or limit their generalization.

i. *Sample size and replication*. Each condition was tested in three wells within a single experiment, without an *a priori* power calculation. Technical replicates capture pipetting and reading variability but not variation between experiments, so the reported intervals probably underestimate the true uncertainty.
ii. *Dosing units*. Concentrations are expressed as % v/v of the purified latex rather than as dry weight (mg/mL), which prevents direct comparison with other studies and between latex batches.
iii. *Chemical characterization*. The extract was not characterized (extraction yield, proanthocyanidin content or chromatographic profile), so the activity cannot be attributed to specific constituents and batch-to-batch reproducibility is unknown.
iv. *Residual solvent and pH*. Residual ethanol was not quantified, no solvent control was included and the pH of the working solutions was not recorded. Residual ethanol or acidity could have contributed to the inhibition and led to overestimation of the latex effect.
v. *Planktonic model and endpoint*. Only planktonic cells were tested. Biofilms on saliva-coated surfaces are generally far less susceptible, and the absence of colony counts leaves bactericidal activity undetermined.
vi. *Strain*. A single reference strain was used, so the results may not extend to clinical isolates.
vii. *Blinding*. Plate preparation and reading were not blinded. The instrumental readout limits, but does not eliminate, the risk of measurement bias.

Future work should replicate these observations in at least three independent experiments, express concentrations in mg/mL of dry extract, characterize the latex chemically, include solvent and pH controls, determine MIC and MBC by standardized broth microdilution [13] with viable counts, and evaluate activity in biofilm models and cytotoxicity toward oral cells.

## 5 Conclusions

Purified *Croton lechleri* latex inhibited the metabolic activity of planktonic *Streptococcus mutans* ATCC 25175 *in vitro* in a concentration-dependent manner, with an estimated IC_50_ of 12.3% v/v (working solution) and near-complete inhibition at the highest concentrations. Given the exploratory design, the small number of technical replicates and the lack of chemical standardization, these results are preliminary. They do not demonstrate equivalence with chlorhexidine and justify confirmatory studies with standardized extracts and biofilm models before any potential role as an oral care adjunct can be assessed.

## Declarations

### Ethics

Not applicable. The study did not involve human participants, human samples or animals.

### Funding

This work received no external funding.

### Competing interests

L.F. Rondán Villagra is a co-founder and minority shareholder of Alveoprol S.A.C.S., a company developing an oral-care hydrogel that contains *C. lechleri* resin. The other authors declare no competing interests.

### Data availability

All replicate-level data are reported in Supplementary Table S1.

## Supplementary material

**Table S1.** Replicate-level data. Metabolic activity relative to the growth control and inhibition for each well.

| Condition | Well | Metabolic activity (%) | Inhibition (%) |
| --- | --- | --- | --- |
| Vehicle (0%) | 1 | 101.987 | -1.987 |
|  | 2 | 99.447 | 0.553 |
|  | 3 | 102.591 | -2.591 |
| 3% | 1 | 99.615 | 0.385 |
|  | 2 | 90.829 | 9.171 |
|  | 3 | 90.829 | 9.171 |
| 6% | 1 | 89.475 | 10.525 |
|  | 2 | 84.605 | 15.395 |
|  | 3 | 77.183 | 22.817 |
| 12% | 1 | 59.340 | 40.660 |
|  | 2 | 51.293 | 48.707 |
|  | 3 | 51.274 | 48.726 |
| 25% | 1 | 36.694 | 63.306 |
|  | 2 | 21.607 | 78.393 |
|  | 3 | 22.926 | 77.074 |
| 50% | 1 | 14.626 | 85.374 |
|  | 2 | 11.923 | 88.077 |
|  | 3 | 19.885 | 80.115 |
| 100% | 1 | 5.460 | 94.540 |
|  | 2 | 2.938 | 97.062 |
|  | 3 | 17.328 | 82.672 |
| Chlorhexidine 1% | 1 | 4.097 | 95.903 |
|  | 2 | 5.270 | 94.730 |
|  | 3 | 0.000 | 100.000 |

**Table S2.** All pairwise comparisons (Tukey HSD). Difference = mean activity of the first condition minus the second, in percentage points, with simultaneous 95% CI.

| Condition 1 | Condition 2 | Difference | 95% CI | p |
| --- | --- | --- | --- | --- |
| Vehicle (0%) | 3% | 7.6 | -7.9 to 23.1 | 0.693 |
| Vehicle (0%) | 6% | 17.6 | 2.1 to 33.1 | 0.021 |
| Vehicle (0%) | 12% | 47.4 | 31.8 to 62.9 | <0.001 |
| Vehicle (0%) | 25% | 74.3 | 58.7 to 89.8 | <0.001 |
| Vehicle (0%) | 50% | 85.9 | 70.3 to 101.4 | <0.001 |
| Vehicle (0%) | 100% | 92.8 | 77.2 to 108.3 | <0.001 |
| Vehicle (0%) | Chlorhexidine 1% | 98.2 | 82.7 to 113.7 | <0.001 |
| 3% | 6% | 10.0 | -5.5 to 25.5 | 0.385 |
| 3% | 12% | 39.8 | 24.3 to 55.3 | <0.001 |
| 3% | 25% | 66.7 | 51.2 to 82.2 | <0.001 |
| 3% | 50% | 78.3 | 62.8 to 93.8 | <0.001 |
| 3% | 100% | 85.2 | 69.7 to 100.7 | <0.001 |
| 3% | Chlorhexidine 1% | 90.6 | 75.1 to 106.2 | <0.001 |
| 6% | 12% | 29.8 | 14.3 to 45.3 | <0.001 |
| 6% | 25% | 56.7 | 41.2 to 72.2 | <0.001 |
| 6% | 50% | 68.3 | 52.7 to 83.8 | <0.001 |
| 6% | 100% | 75.2 | 59.7 to 90.7 | <0.001 |
| 6% | Chlorhexidine 1% | 80.6 | 65.1 to 96.2 | <0.001 |
| 12% | 25% | 26.9 | 11.4 to 42.4 | <0.001 |
| 12% | 50% | 38.5 | 23.0 to 54.0 | <0.001 |
| 12% | 100% | 45.4 | 29.9 to 60.9 | <0.001 |
| 12% | Chlorhexidine 1% | 50.8 | 35.3 to 66.4 | <0.001 |
| 25% | 50% | 11.6 | -3.9 to 27.1 | 0.230 |
| 25% | 100% | 18.5 | 3.0 to 34.0 | 0.014 |
| 25% | Chlorhexidine 1% | 24.0 | 8.4 to 39.5 | 0.001 |
| 50% | 100% | 6.9 | -8.6 to 22.4 | 0.777 |
| 50% | Chlorhexidine 1% | 12.4 | -3.2 to 27.9 | 0.175 |
| 100% | Chlorhexidine 1% | 5.5 | -10.1 to 21.0 | 0.916 |

## Notes

### Competing Interest Statement

L.F. Rondan Villagra is a co-founder and minority shareholder of Alveoprol S.A.C.S., a company developing an oral-care hydrogel that contains Croton lechleri resin. The other authors declare no competing interests.

